# The inhibitory effects of butein on cell proliferation and TNF-α-induced CCL2 release in racially different triple negative breast cancer cells

**DOI:** 10.1101/596080

**Authors:** Patricia Mendonca, Ainsley Horton, David Bauer, Samia Messeha, Karam F.A. Soliman

## Abstract

Breast cancer drug resistance is the leading cause of cancer-related mortality in women, and triple negative breast cancer (TNBC) is the most aggressive subtype, affecting African American women more aggressively compared to Caucasians. Of all cancer-related deaths, 15 to 20% are associated with inflammation, where proinflammatory cytokines have been implicated in the tumorigenesis process. The current study investigated the effects of the polyphenolic compound butein (2′,3,4,4′-tetrahydroxychalcone) in cell proliferation and survival, as well as its modulatory effect on the release of proinflammatory cytokines in MDA-MB-231 (Caucasian) and MDA-MB-468 (African American) TNBC cell. Results showed that butein decreased cell viability in a time and dose-dependent manner and after 72-h of treatment, cell proliferation rate was reduced in both cell lines. In addition, butein presented higher potency in MDA-MB-468, exhibiting anti-proliferative effects in lower concentrations. Apoptosis assays demonstrated that butein increased apoptotic cells in MDA MB-468, showing 90% of the analyzed cells in the apoptotic phase, compared to 54% in MDA-MB-231 cells. Additionally, butein downregulated both, protein and mRNA expression of CCL2 proinflammatory cytokine and IKBKE in Caucasian cells, but not in African Americans. This study demonstrates butein potential in cancer suppression showing a higher cytotoxic, anti-proliferative, and apoptotic effects in African Americans, compared to Caucasians TNBC cells. It also reveals the butein inhibitory effect on CCL2 expression with a possible association with IKBKE downregulation in MDA-MB-231 cells only, indicating that Caucasians and African Americans TNBC cells respond differently to butein treatment. The obtained findings may provide an explanation regarding the poor response to therapy in African American patients with advance TNBC.

## Introduction

Natural compounds isolated from medicinal plants have been the source of novel agents that stimulate numerous physiological pathways and can be beneficial to prevent and cure many diseases, including cancer [1–3]. For decades, new approaches to investigate the effects of phytochemicals with promising therapeutic potential and reduced adverse side effects have been explored [4–6]. Butein (2',3,4,4'-tetrahydroxychalcone) is a polyphenol compound found in several plants, including *Semecarpus anacardium*, *Dalbergia odorifera*, and *Rhus verniciflua* Stokes [7]. In Asian countries, butein has been used in herbal medicine formulations and as a food additive [8]. Also, butein exhibits a variety of pharmacological properties including anti-inflammatory, antioxidative, and antimicrobial activities [9,10].

In cancer studies, butein induced apoptosis in colon adenocarcinoma [11], breast cancer cells [12,13], hepatocellular carcinoma [14,15], and prostate cancer cells [16,17]. Butein treatment also inhibited invasion and migration of bladder [18], pancreatic, and breast cancer cells [19], as well as in hepatocellular carcinoma [20]. Breast cancer cell studies showed that butein inhibited ER^+^ MCF-7 cells growth [12], and blocked CXCL12-induced migration and invasion of human epidermal growth factor receptor 2 positive (HER2^+^) in SKBR-3 breast cancer cells by repressing NFκB-dependent CXCR4 expression [19]. Moreover, the compound induced apoptosis in MDA-MB-231, though ROS generation and ERK1/2 and p38MAPK dysregulation [13]. These findings show butein potential as a promising chemopreventive and chemotherapeutic potential agent [21].

The high prevalence and increasing drug resistance of breast cancer is the leading cause of cancer-related mortality in women [22]. In 2018, there was an estimated number of 266,000 new cases of invasive breast cancer to be diagnosed in the U.S., alongside 64,000 new cases of non-invasive breast cancer [23]. Breast cancer is classified into three major therapeutic subtypes: estrogen and/or progesterone receptor-positive (ER^+^, PR^+^), HER2^+^, and triple-negative breast cancer (TNBC) (lacking expression of ER, PR, and HER2) [24,25]. TNBC covers 15 to 20% of all breast cancers [26], and it is linked to a worse prognosis, increased incidence of metastasis to lung, liver, and brain, early relapse after treatment, and lower survival rate compared to other breast cancer subtypes [27]. TNBC is more common in African American women compared to other ethnic groups [28,29] and it is associated with a worse clinical outcome, and it presents higher mortality [27,30]. TNBC subtypes respond differently to the treatment, challenging, even more, the development of target therapy with certain chemotherapeutics that may be safe and effective at the same time [25,31].

In addition to breast cancer heterogeneity, it is also known that 15 to 20% of all cancer-related deaths worldwide are associated with inflammation [32]. Also, the existence of the relationship between cancer and stromal cells at the tumor site, set up by inflammatory cytokines has been documented [33]. Proinflammatory cytokines are the crucial link between chronic inflammation and cancer, inducing tumor progression, proliferation, proangiogenic factors, inhibiting apoptosis and immune-mediated tumor surveillance, and thus facilitating carcinogenesis [34,35].

Despite the availability of evidence confirming butein effectiveness in tumor suppression, there is meager research information regarding its influence on the tumor cell response to proinflammatory cytokines, specifically TNF-α. In BC, high concentrations of TNF-α can activate receptors and trigger a potent and persistent activation of NFκB signaling [33,36], epithelial-to-mesenchymal transition [37], and continuous release of diverse chemokines, including CCL2 and CCL5 [38]. These chemokines may initiate an inward migration of numerous leukocyte sub-populations (LPSs), including tumor-associated macrophages[39], myeloid-derived suppressor cells [40], tumor-associated neutrophils [41,42], T-regulatory [43], metastasis-associated macrophages, T helper IL-17-producing cells, and cancer-associated fibroblasts [44], which may bear CCR2 / CCR5 receptors, driving tumor aggression [45,46]. Therefore, chemokines are recognized as key trafficking molecules produced by cancer cells in response to TNF-α stimulation, and able of driving LSPs recruitment [47–50].

Although evidence in the literature show butein potential in protecting against and suppressing cancer, there are no studies to compare the effect of this compound on TNF-α-induced CCL2 release in Caucasian and African American breast cancer cell lines. Therefore, the present work is investigating the effect of the polyphenol compound butein in different TNBC cells. Butein effects on cell viability and cell proliferation were evaluated, as well as butein modulatory effect in the release of TNF-α-induced proinflammatory cytokines.

## Materials and Methods

### Cell lines, Chemicals, and Reagents

MDA-MB-231 (Caucasian American) and MDA-MB-468 (African American) TNBC cells were purchased from American Type Culture Collection (ATCC). Dulbecco’s modified Eagle’s medium (DMEM) high glucose; fetal bovine serum heat inactivated (FBS-HI), penicillin/streptomycin and Hank's Balanced salt solution (HBSS) were obtained from Genesee Scientific (San Diego, CA, USA). Dimethyl sulfoxide (DMSO), butein, and Alamar Blue^®^ were purchased from Sigma-Aldrich Co. (St. Louis, MO, USA). Human cytokine antibody arrays (Cat# AAH-CYT-6-4), ELISA assays for MCP-1 (Cat# ELH-MCP1-1), Annexin V-FITC apoptosis Kit (Cat# 68FT-AnnV-S100), and tumor necrosis factor alpha (TNF-α) were purchased from RayBiotech (Norcross, Ga, USA). PCR primers and iScript advanced reverse transcriptase kit were purchased from Bio-Rad (Hercules, CA, USA). DNA-free™ Kit (Cat # AM1907) was obtained from Life Technologies Inc. (Grand Island, NY, USA).

### Cell culture

MDA-MB-231 and MDA-MB-468 TNBC cells were cultured in DMEM supplemented with 10% FBS-HI and 1% penicillin (100 U/ml)/streptomycin (0.1 mg/ml) and incubated in an atmosphere of 5% CO_2_ and 37°C. Cells were sub-cultured in T-75 flasks and grown to 90% confluency before setting the cells for each assay. Plating media for each experiment consisted of DMEM, with 2.5% of FBS-HI, and no penicillin/streptomycin.

### Cell viability

Alamar Blue^®^ (Resazurin) assay was used to assess MDA-MB-231 and MDA-MB-468 cell viability. Briefly, 96-well plates were seeded with cells at a density of 3×10^4^ cells/100 μl/well and incubated overnight to attach. The next day, the cells were treated as follows: control (media only), control (cells + DMSO), and cells treated with different concentrations of butein (0.78 - 200 μM). Butein was dissolved in DMSO before dilution in the media and the final concentration of DMSO did not exceed 0.1%. The volume of 100 μl of each treatment was added to the plate-containing cells. The butein effect was measured after different periods of incubation: 24, 48, and 72 h. The amount of 20 μl of Alamar Blue^®^ solution (0.5 mg/ml) was added to the plate and incubated again for 4 h. Quantitative analysis of dye conversion was measured at an excitation/emission of 550/580 nm wavelengths using a microplate reader Infinite M200 (Tecan Trading AG). Viable cells were able to reduce resazurin to resorufin, resulting in fluorescence changes. The fluorescent signal was proportional to the number of living cells in the sample, and the data were expressed as a percentage of alive untreated controls.

### Cell proliferation assay

Butein effect on cell proliferation was determined in MDA-MB-231 and MDA-MB-468 TNBC cell lines based on the cell viability study concentrations by using Alamar Blue^®^. Cells were plated at an initial density of 5×10^3^ cell/well in 96-well plates and incubated overnight in experimental media. The next day, the cells were treated with butein at concentrations ranging from 0.78 - 200 μM, in a final volume of 200 μl/well. Control cells were exposed to DMSO at a concentration of < 0.1%, as described above. Taxol (1 μM) was used as a positive control. Proliferation was measured after 72 h of butein treatment by adding 20 μl of Alamar Blue^®^ and reading the plate at an excitation/emission of 550/580 nm wavelengths using a microplate reader Infinite M200 (Tecan Trading AG). Cell proliferation was calculated based on the percentage of cell growth observed in the control samples.

### Apoptosis assay

The effect of butein in inducing apoptosis was determined in MDA-MB-231 and MDA-MB-468 cells by using Annexin V-FITC Apoptosis assay Kit from RayBiotech. Briefly, each cell line was seeded at an initial concentration of 5×10^5^ cell/well in 6-well plates and incubated overnight. Cells were treated with butein at concentrations ranging between 0 - 200 μM in a final volume of 3 ml/well of experimental media to induce apoptosis. Control cells were exposed to DMSO at a concentration < 0.1%. After 24 h incubation period, controls and treated cells from each well were harvested, pelleted, and washed with PBS. According to the manufacture’s protocol, the cell pellets were resuspended in 500 μl of 1X Annexin -V binding buffer, then labeled with 5 μl of Annexin V-FITC, and 5 μl propidium iodide. The apoptotic effect was quantified within 5 - 10 min by FACSCalibur Flow cytometer (Becton Dickinson, San Jose, CA, USA). For each sample, 1 × 10^4^ cells were examined, and CELLQuest software was used for the data analysis.

### Human cytokine antibody array membrane

RayBiotech human cytokine antibody arrays were used to study the effect of butein on 60 cytokine proteins released by TNF-α-activated TNBC cells. Each experiment was performed in triplicate and according to the manufacturer’s instructions. Shortly, antibody-coated array membranes were first incubated for 30 min with 1 ml of blocking buffer. Then, blocking buffer was decanted and replaced with 1 ml supernatant from control (cells + DMSO) samples, cells treated with butein (5 μM), TNF-α (40 ng/ml), and the combination of butein (5 μM) + TNF-α (40 ng/ml). Membranes were incubated overnight at 4°C with mild shaking. The next day, the media were decanted; membranes were washed, and subsequently incubated with 1 ml biotin-conjugated antibodies for 2 h. Lastly, biotin-conjugated antibodies were removed, and membranes were washed again and incubated with HRP-conjugated streptavidin for 2 h. In this assay chemiluminescent reagent was used and the image of spots was captured using a Flour-S Max Multi-imager (Bio-Rad Laboratories, Hercules, CA, USA), and the spot density was determined with Quantity One Software (Bio-Rad Laboratories, Hercules, CA). Excel-based data analysis was performed, using Human Cytokine Array software C1000 (CODE: S02-AAH-CYT-1000) from RayBiotech.

### Human CCL2 (MCP-1) ELISA quantification

Supernatants from the control (cells + DMSO), butein-treated, TNF-α-stimulated, and co-treated (butein (5 μM) + TNF-α (40 ng/ml) TNBC cells were collected and centrifuged at 1000 rpm for 4 min at 4°C. Specific ELISA assays for CCL2 (MCP-1) was performed following the manufacturer’s instructions. Shortly, 100 μl of supernatants from each sample and standards were added to 96 well plates pre-coated with capture antibody and incubated for 2.5 h at room temperature under shaking. After washing, 100 μl of prepared biotinylated antibody mixture was added to each well and incubated for 1 h. The mixture was decanted, and 100 μl streptavidin solution was added to each well and incubated for 45 min. Substrate reagent (100 μl) was then pipetted into each well and incubated for 30 min, followed by the addition of 50 μl of stop solution. Data were quantified by optical density at 450 nm using Synergy HTX Multi-Reader (BioTek, USA).

### Real time polymerase chain reaction (RT-PCR)

#### RNA extraction

Cell pellets from control (cells + DMSO), butein-treated (5 μM), TNF-α-stimulated (40 ng/ml) and co-treated with butein (5 μM) + TNF-α (40 ng/ml) were lysed with 1ml TRIzol reagent. Then, chloroform (0.2 ml) was added to the lysed samples; the tubes were shaken, incubated at 15 - 30°C for 2 - 3 min, and centrifuged at 10,000 rpm for 15 min at 2 - 8°C. Lysed samples (aqueous phase) were then transferred to a new tube, and mixed with 0.5 ml of isopropyl alcohol for RNA precipitation. After incubation (15 min), samples were centrifuged, the supernatant was removed, the RNA pellets were washed with 75% ethanol (by inverting the tubes carefully), and then centrifuged at 7,500 rpm for 5 min at 2 - 8°C. The RNA pellet was dried (room temperature), dissolved in RNase-free water, and incubated on ice (30 min). Finally, using Nanodrop (Thermo Fischer Scientific, Wilmington, DE, USA), RNA purity and quantity were determined.

#### cDNA synthesis and RT-PCR

The cDNA strands were synthesized from the mRNA using iScript advanced reverse transcriptase from Bio-Rad. A solution of 4 μl of the 5X iScript advanced reaction mix (containing primers), 1 μl of reverse transcriptase, 7.5 μl of the sample (1.5 μg/reaction), and 7.5 μl of water were combined in a 0.2 ml tubes, in a total volume of 20 μl. The thermal cycling program for the reverse transcription included two steps: 46°C for 20 min and then 95°C for 1 min. RT-PCR amplification was performed following the manufacturer protocol (Bio-Rad). A 1 μl of the sample (200 ng cDNA/reaction), 10μl of master mix, 1 μl of primer, and 8 μl of water were combined into each well. The thermal cycling process included an initial hold step at 95°C for 2 min and denaturation at 95°C for 10 sec, followed by 39 cycles of 60°C for 30 sec (annealing/extension), and 65°C - 95°C for 5 sec/step (melting curve) using the Bio-Rad CFX96 Real-Time System (Hercules, CA, USA). The selected primers were specific to each gene of interest. The UniqueAssay ID for *CCL2/MCP1* primer was qHsaCID0011608, and the for *IKBKE* primer was qHsaCID0014831.

### Statistical Analysis

Data analysis was performed using GraphPad Prism (version 6.07) (San Diego, CA, USA). All data points are expressed as the mean ± S.E.M. from at least 2 independent experiments. For the viability studies, the IC_50_ was determined by nonlinear regression with R^2^ best fit and lowest 95% confidence interval. Statistically significant differences between different groups in the experiments was assessed using a one-way ANOVA, followed by Dunnett’s multiple comparison tests (*P < 0.05, **P < 0.01, ***P < 0.001, ****P < 0.0001, and ns= p > 0.05). Gene expression was analyzed using the CFX 3.1 Manager software (Bio-Rad, Hercules, CA).

## Results

Butein effect on breast cancer cell viability was investigated in MDA-MB-231 and MDA-MB-468 cell lines after 24 h-treatment. Butein caused a concentration-dependent decrease in the cell viability in both cell lines. Low concentrations of butein (from 0.78 to 6.25 μM) showed no cytotoxicity in MDA-MD-231; however, in MDA-MD-468, the decrease in cell viability was statistically significant (p < 0.0001) in the lowest concentration of 0.78 μM, compared to the control. The IC_50_ of 54.07 ± 2.8 μM for Caucasian and 33.75 ± 1.6 μM for African American indicate that butein effects on the two cell lines are different, causing higher cytotoxicity effect in MDA-MB-468 cells (**Fig 1**).

**Fig 1:**
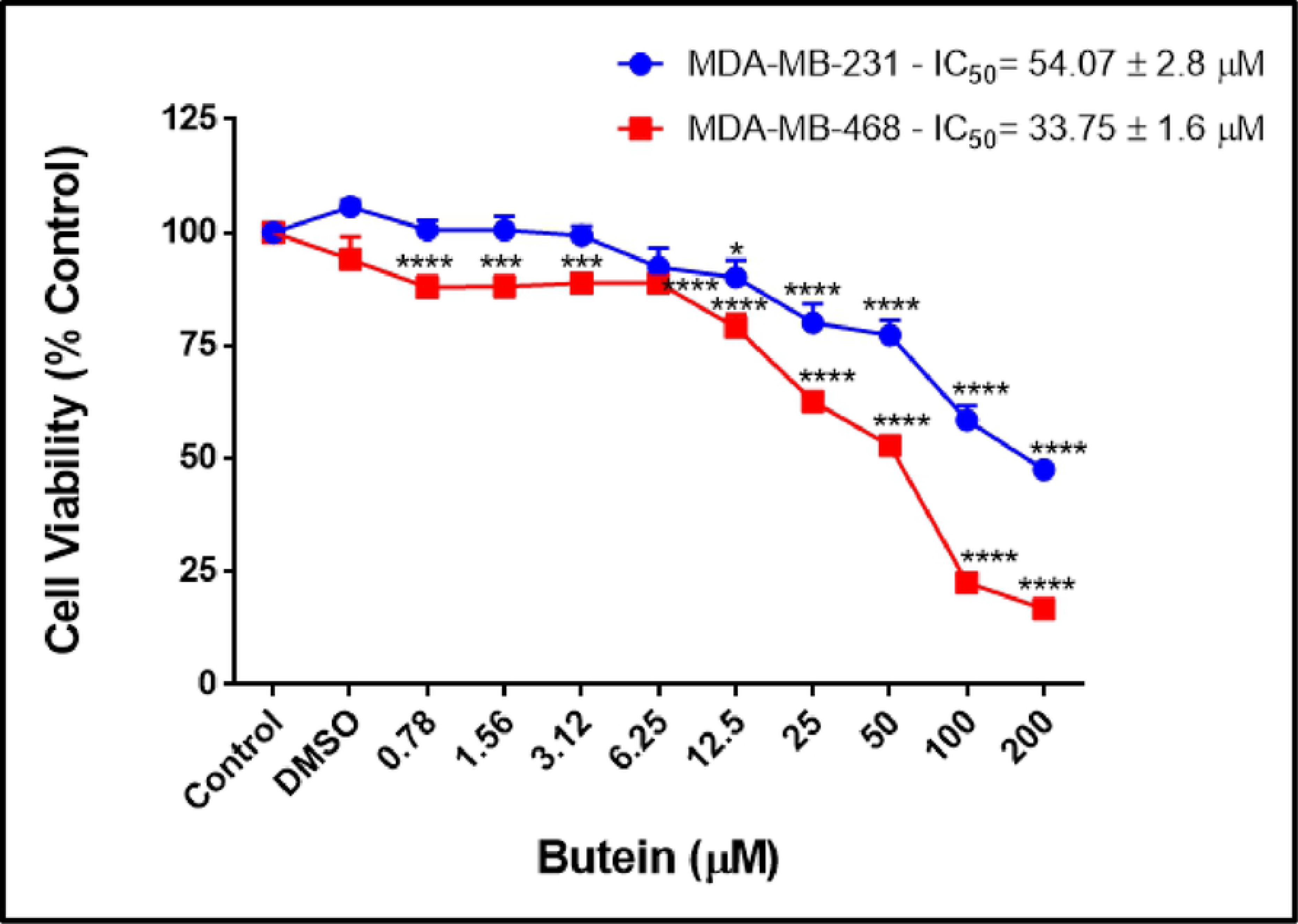
The effect of butein on cell viability in MDA MB-231 and MDA MB-468 TNBC cells after the 24-h treatment period. Butein concentrations ranged from 0.78 - 200 μM. All experiments were performed at least 3 times with n=5 and kept for 24 h at 5% CO_2_ and 37°C. The data are presented as the mean ± S.E.M. Statistically significant differences between groups (control vs. treatments) were evaluated by a one-way ANOVA, followed by Dunnett’s multiple comparison tests. *p < 0.05, ***p < 0.001, ****p < 0.0001.

The cytotoxic effect of butein was examined by incubating both cell lines with butein for 24, 48 and 72 h. Results obtained indicate that cell viability rate was inversely correlated with the butein concentrations and exposure periods. At the 48-h incubation period, butein decreased the IC_50_s from 54.07 to 5.82 μM in MDA-MB-231, and from 33.75 to 8.70 μM in MDA-MB-468 cells. Further decrease in the cell viability was also measured at 72-h incubation period, reducing IC_50_s to 1.11 and 1.78 μM in MDA-MB-231 and MDA-MB-468, respectively (**Figs 2A and B**).

**Fig 2:**
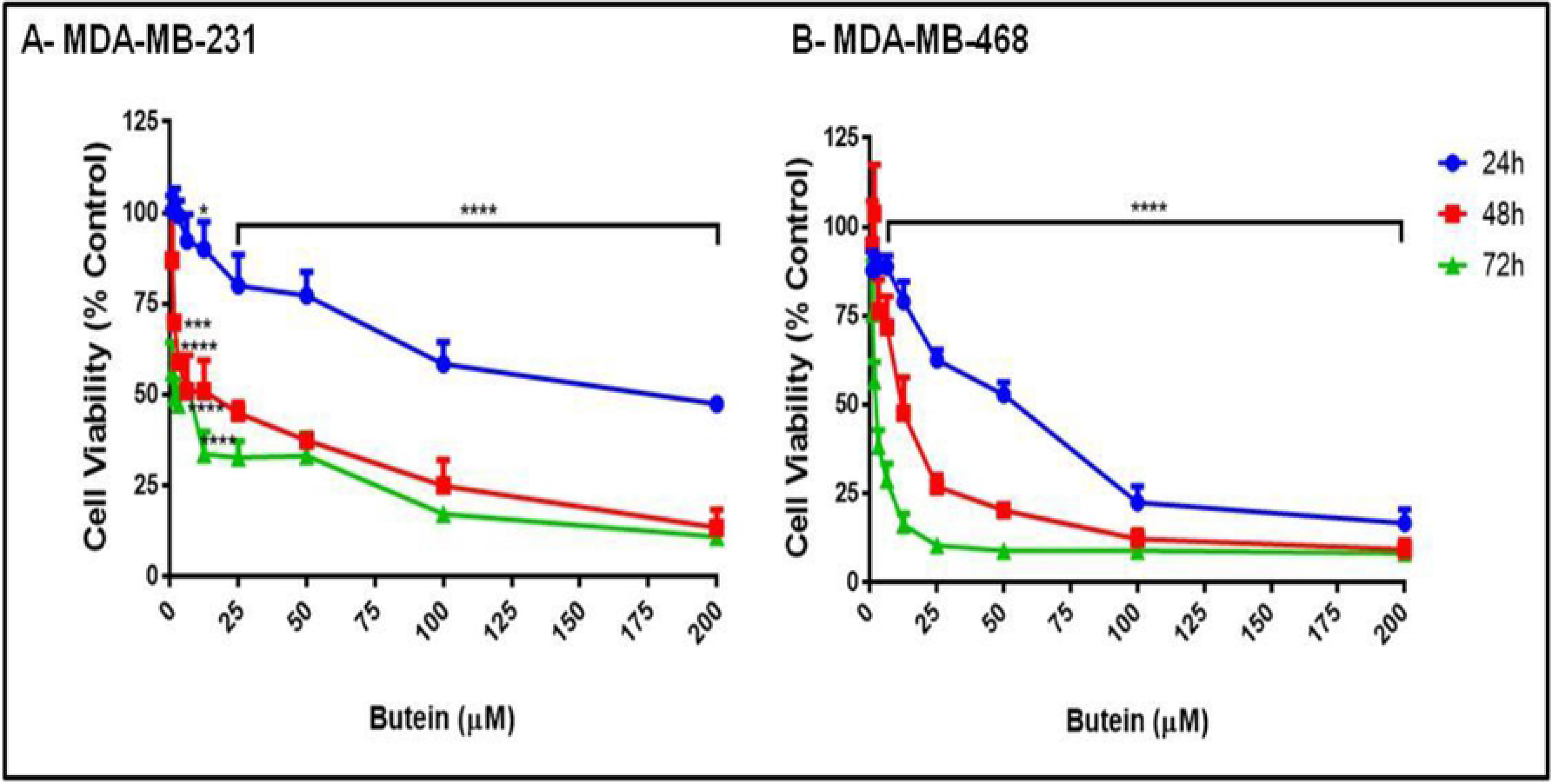
The cytotoxic effect of butein in (A) MDA MB-231 and (B) MDA MB-468 TNBC cells at 24, 48, and 72-h treatment period. Each cell line was treated up to 72 h with butein at concentrations ranging from 0.78 - 200 μM. All experiments were performed at least 3 times with n=5 and kept at 5% CO_2_ and 37°C. Each data point represent the mean ± S.E.M. Statistically significant differences between groups (control vs. treatments) were evaluated by a one-way ANOVA, followed by Dunnett’s multiple comparison tests. *p < 0.05, ***p < 0.001, ****p < 0.0001.

Anti-proliferation assays, based on the resazurin reduction, were performed to determine the potency of butein in inhibiting cell growth of both cell lines. The cells were treated with butein at concentrations ranging from 0.78 to 200 μM. The decrease in cell proliferation rate was detected in a dose-dependent manner. In MDA-MB-468, butein started exerting its effect in a lower concentration (6.25 μM), compared to MDA-MB-231 cells (12.5 μM). Similar to the chemotherapy drug Taxol after the 72-h treatment period, butein reduced breast cancer cells growth and showed its potency as an anti-proliferative agent (**Figs 3A and B**).

**Fig 3:**
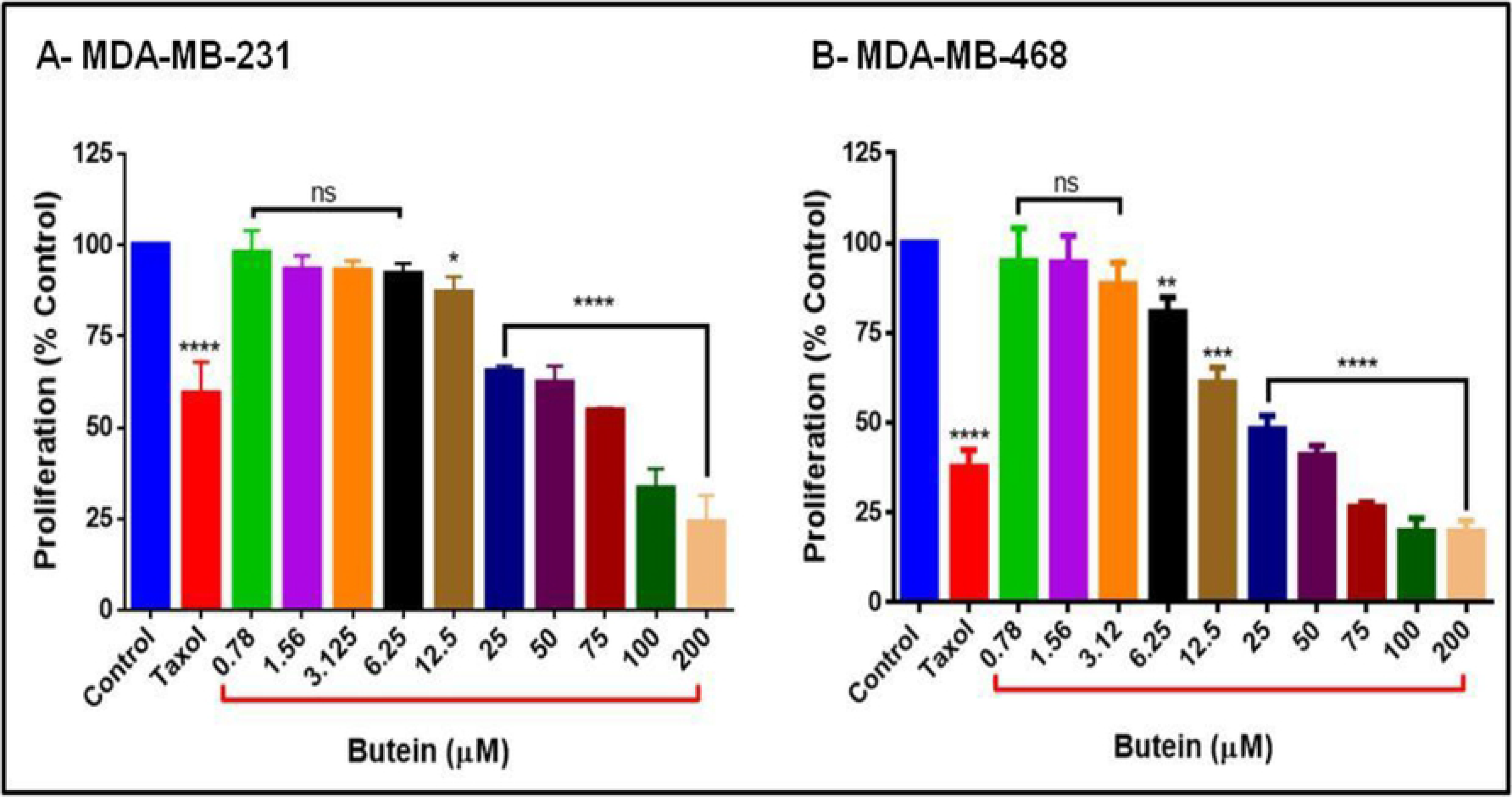
The effect of butein on cell proliferation in (A) MDA MB-231 and (B) MDA MB-468 TNBC cells. Butein concentrations ranging from 0.78 - 200 μM and Taxol (1 μM) were tested in both cell lines and kept at 5% CO_2_ and 37°C for a 72-h incubation period. Each data point represents the mean ± S.E.M. of four independent experiments (n=5). Statistically significant differences between groups (control vs treatments) were evaluated by a one-way ANOVA, followed by Dunnett’s multiple comparison tests. *p < 0.05, **p < 0.01, ***p < 0.001, ****p < 0.0001, ns= p > 0.05.

The apoptotic effect of butein was determined by flow cytometry using Annexin V-FITC/PI staining in cells exposed to butein for 24 h. In MDA-MB-231 cells, there was a significant increase of early apoptosis and a progressive increase of late apoptosis with increasing concentrations of butein. Treatment with concentrations of 12.5 (lowest) and 200 μM (highest) of butein increased the percentage of apoptotic cells (early and late) from 12.02 ± 1.8% to 54.0 ± 0.37 (**Figs 4A and B**). Moreover, MDA-MB-468 cells were more sensitive to butein compared to MDA-MB-231, presenting apoptosis in 90.0 ± 1.44 % of the cells at the highest concentration of butein (200 μM) after 24-h treatment (**Figs 5A and B**). These results demonstrate that butein was more effective in inducing apoptosis in African American compared to Caucasians in TNBC cells.

**Fig 4:**
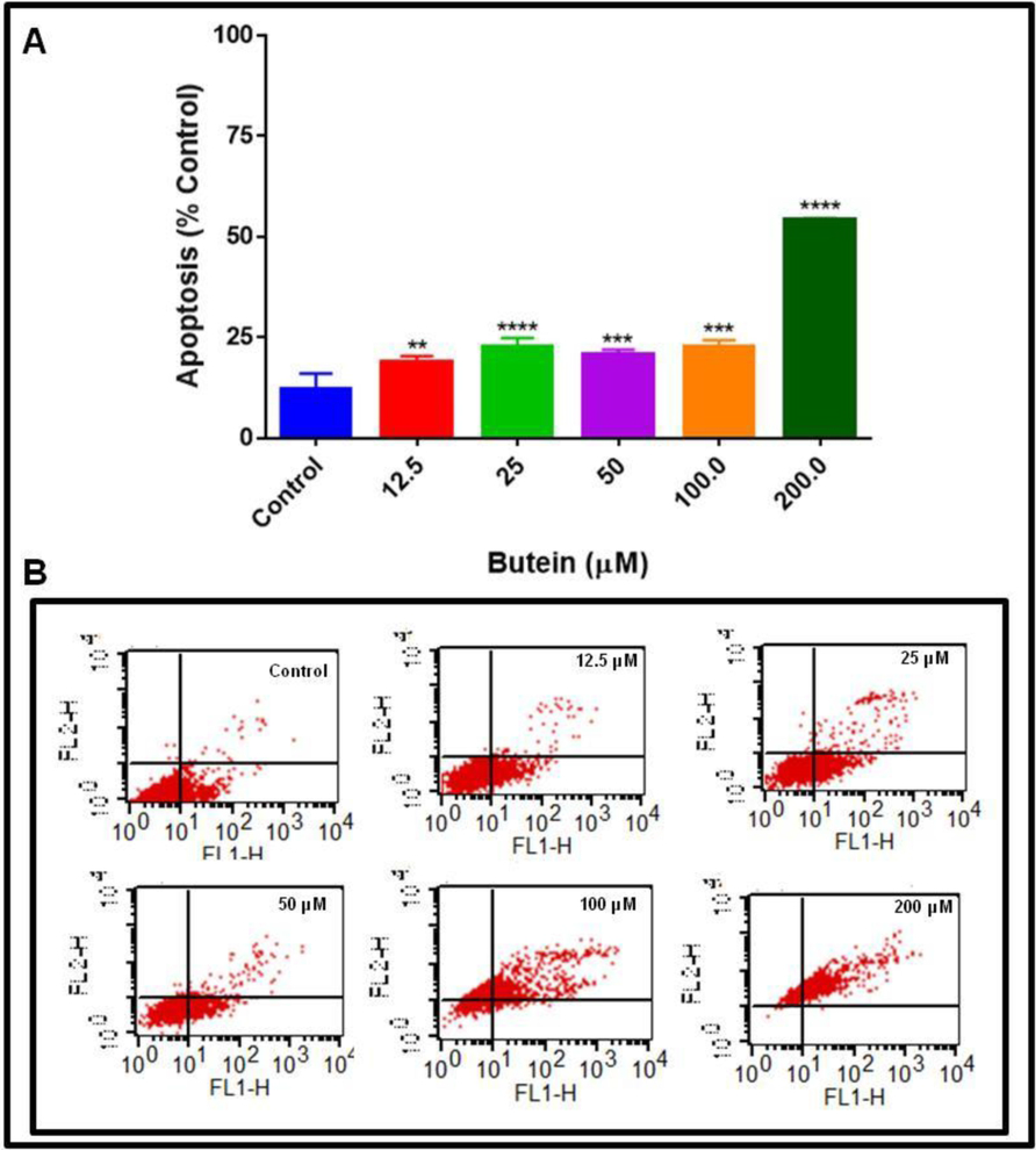
The apoptotic effect of butein in MDA-MB-231 TNBC cell line. Cells were exposed to butein at concentrations ranging from 12.5 - 200 μM for 24 h, and control cells were treated with DMSO (< 0.1%). Apoptotic effect was determined by flow cytometry using Annexin V-FITC kit and FACSCalibur Flow cytometer to analyze the percentage of the apoptotic cells compared to the control cells. The results represent the mean ± S.E.M. of two independent studies (n=3). Statistically significant differences between groups (control vs. treatments) were evaluated by a one-way ANOVA, followed by Dunnett’s multiple comparison tests. **p < 0.01, ***p < 0.001, ****p < 0.0001.

**Fig 5:**
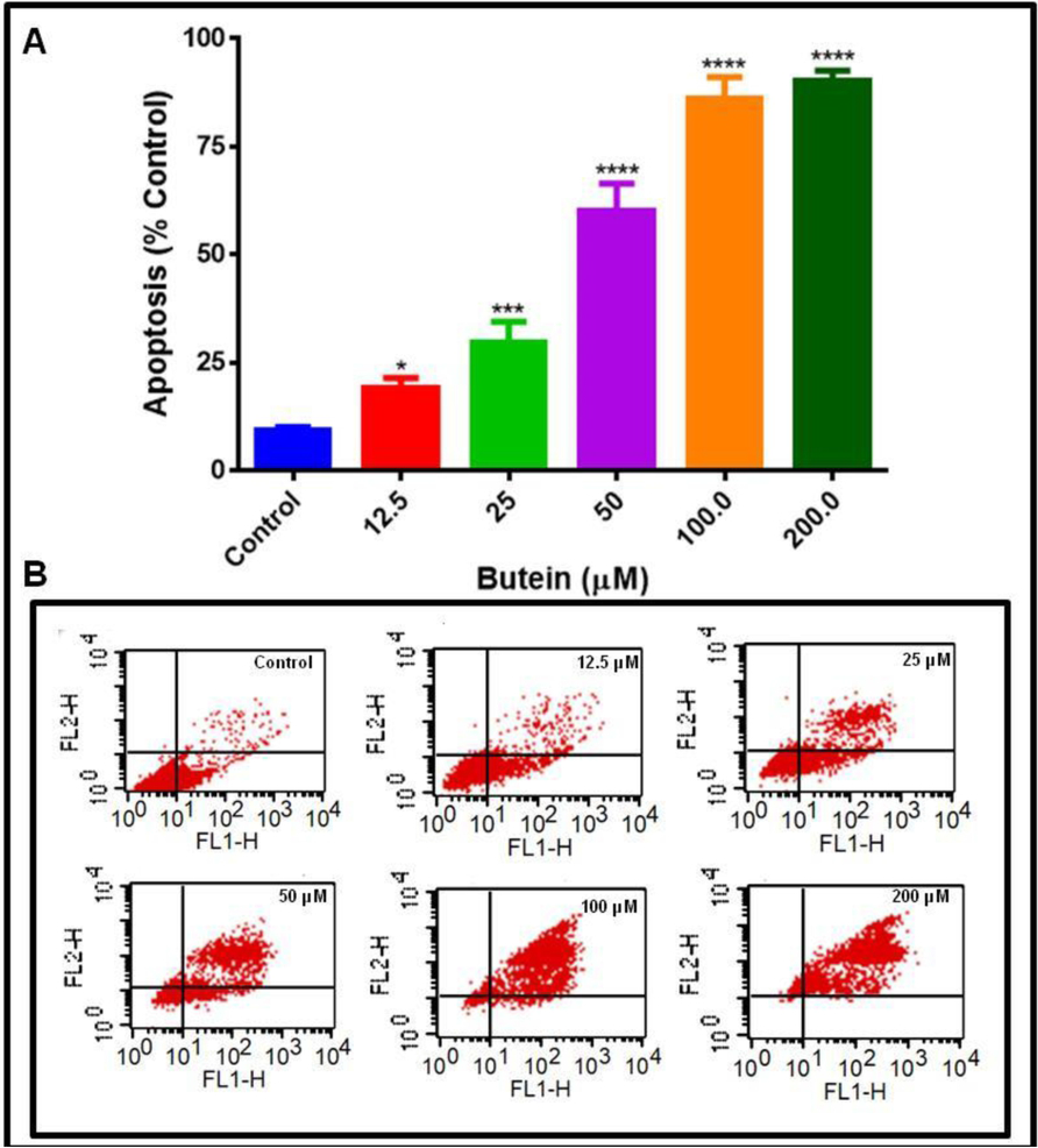
The apoptotic effect of butein in MDA-MB-468 TNBC cell line. Cells were exposed to butein at concentrations ranging from 12.5 - 200 μM for 24 h. Control were treated with DMSO. The apoptotic effect of butein was determined by flow cytometry using Annexin V-FITC kit and FACSCalibur Flow cytometer to analyze the percentage of the apoptotic cells compared to the control cells. The results represent the mean ± S.E.M. of two independent studies (n=3). Statistically significant differences between groups (control vs. treatments) were evaluated by a one-way ANOVA, followed by Dunnett’s multiple

In order to evaluate the relationship between the anti-cancer effects of butein treatment and the inhibitory effect on pro-cytokines release, a semi-quantitative analysis using human antibody arrays was performed (**Figs 6A and 7A**). The results showed that TNF-α induced the upregulation of three specific cytokines: chemokine (C-C motif) ligand 2 (CCL2/MCP-1), insulin-like growth factor-binding protein 1 (IGFBP1), and interleukin-6 (IL-6) in MDA-MB-231 cells, although CCL2 was the only one upregulated in its counterpart MDA-MB-468 (**Figs 6B and 7B**). Butein presented a different effect in the two cell lines examined, inhibiting CCL2 expression in Caucasian, but not in African American cells. A dot blot intensity analysis of the arrays was performed using Quantity One software (Bio-Rad), where the dots intensities were normalized as a percentage of positive controls for each membrane. The results obtained from TNF-α-stimulated cells and cells co-treated with butein + TNF-α show that butein attenuated TNF-α-induced CCL2 release significantly in MDA-MB-231 (4-fold inhibition), but not in MDA-MB-468 (**Figs 8A and B**).

**Fig 6:**
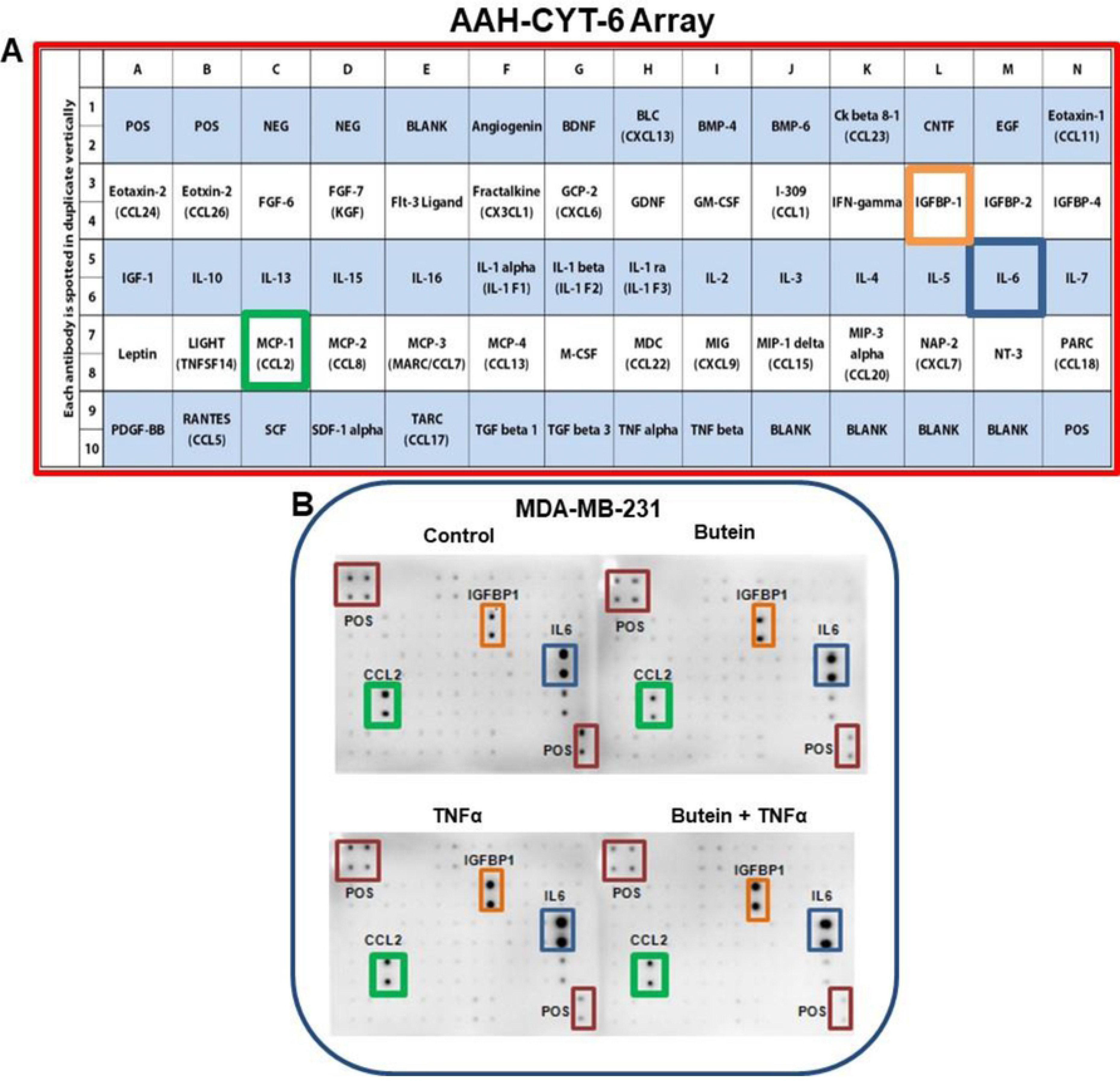
The effect of butein on cytokine expression in TNF-α-activated MDA-MB-231 TNBC cells (n=3). **A-** Array layout used to assess chemokines/cytokines expression in the cell-free supernatants, showing the cytokines map, and highlighting CCL2 (MCP1), IGFBP-1, and IL-6. **B -** Array with chemiluminescent spot intensity analysis of supernatants derived from Caucasian and African American breast cancer cells showing cytokine changed expression after treatments. Blots represent the supernatants of 4 treatments: control (cells + DMSO), butein (5 μM), TNF-α (40 ng/ml), and butein (5 μM) + TNF-α (40 ng/ml).

**Fig 7:**
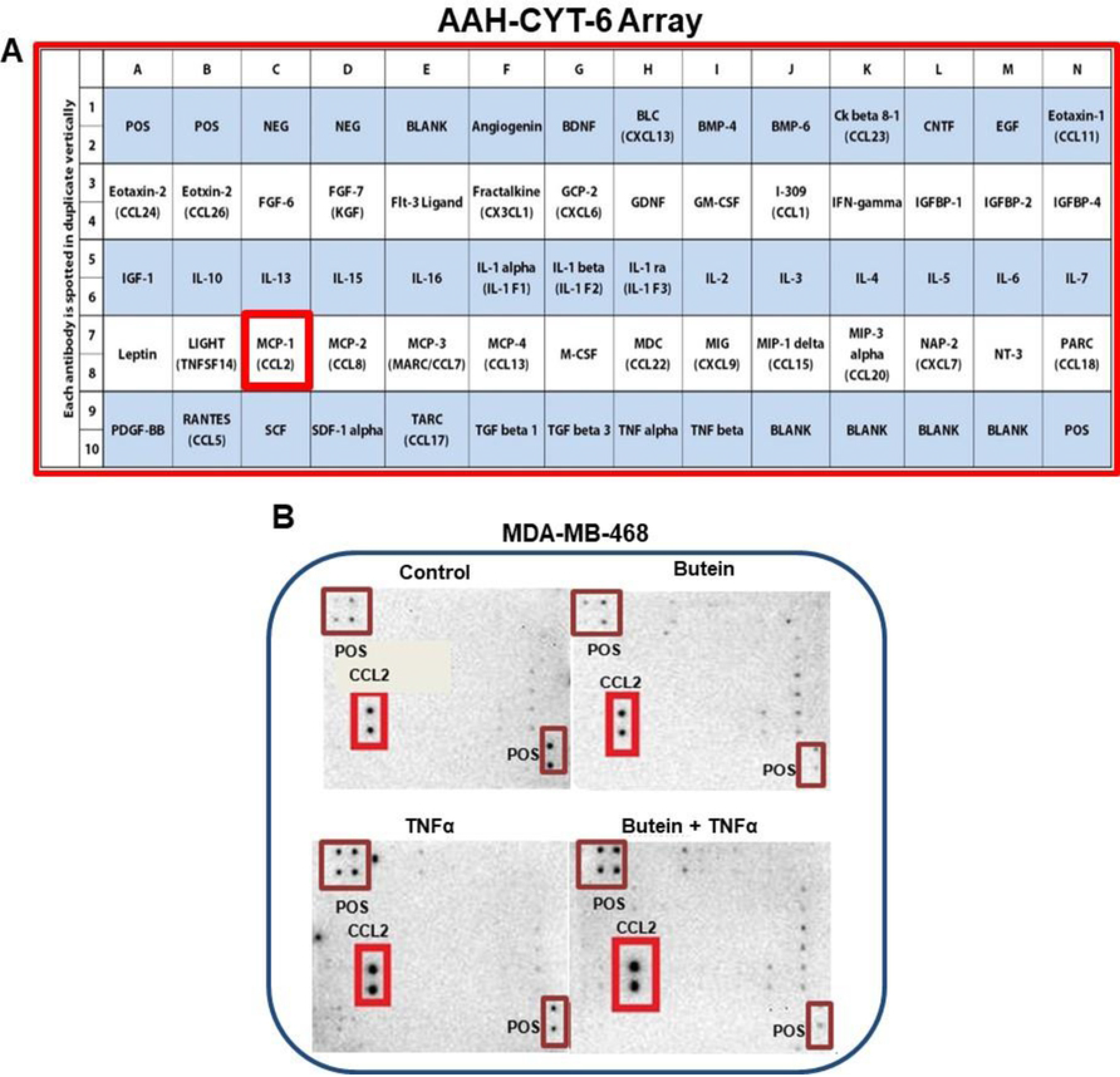
The effect of butein on cytokine expression in TNF-α-activated MDA-MB-468 TNBC cells (n=3). **A-** Array layout used to assess chemokines/cytokines expression in the cell-free supernatants, showing the cytokines map, and highlighting CCL2 (MCP1). **B -** Array with chemiluminescent spot intensity analysis of supernatants derived from Caucasian and African American breast cancer cells showing cytokine changed expression after treatments. Blots represent the supernatants of 4 treatments: control (cells + DMSO), butein (5 μM), TNF-α (40 ng/ml), and butein (5 μM) + TNF-α (40 ng/ml).

**Fig 8:**
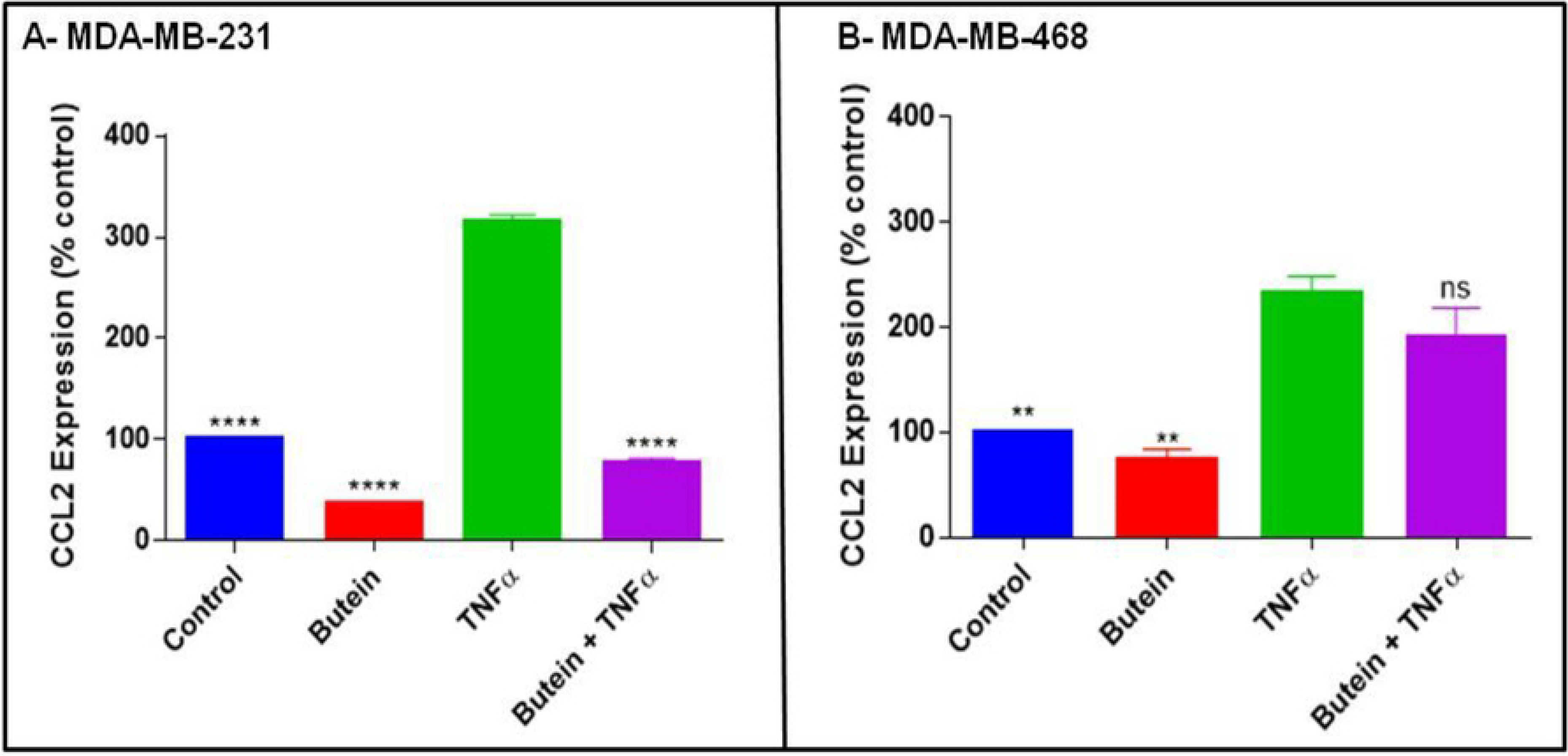
Protein expression of CCL2 in (A) MDA MB-231 and (B) MDA MB-468 TNBC cells. Data are presented as normalized intensities, expressed as % of control arrays (mean ± S.E.M. n= 3). Statistically significant differences between TNF-α vs other treatments were evaluated by a one-way ANOVA, followed by Dunnett’s multiple comparison tests. **p < 0.01, ****p < 0.0001, ns= p > 0.05.

ELISA quantitative assays specific for CCL2 were used to validate the cytokine array findings. The results confirmed that TNF-α induces up-regulation of CCL2 expression in both breast cancer cell lines. However, butein was able to downregulate the same cytokine only in MDA-MB-231, with no significant effect on MDA-MB-468 cells. These findings corroborate butein effect using the cytokine arrays (**Figs 9A and B**).

**Fig 9:**
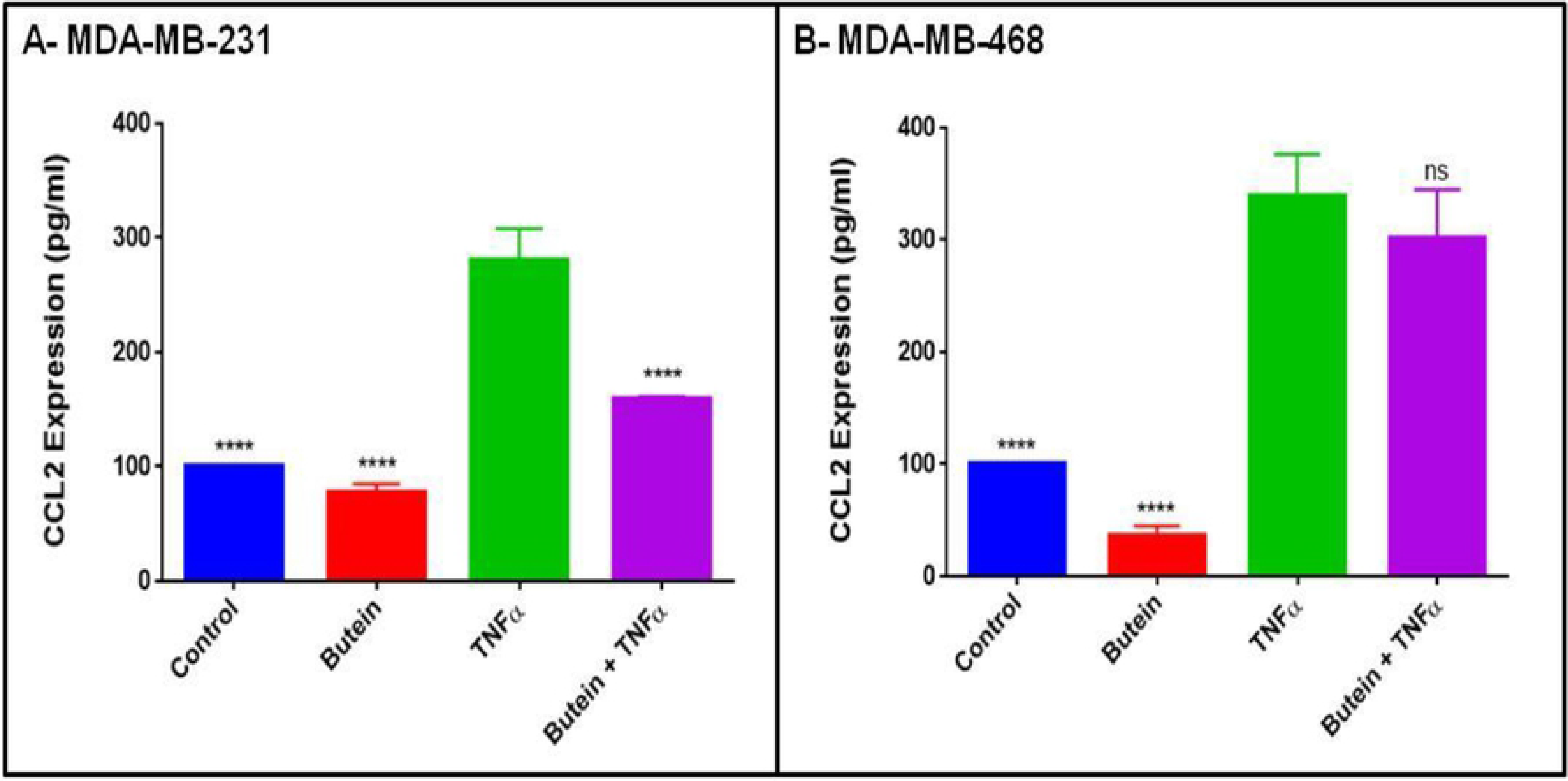
ELISA assay for CCL2 quantification in TNBC cell lines. The effect of butein (5 μM) on CCL2 (MCP1) expression in TNF-α stimulated MDA MB-231 (**A**) and MDA MB-468 cells (**B**). Each data point represents the mean ± S.E.M. of three independent experiments (n=3). Statistically significant differences between TNF-α vs. other treatments were evaluated by a one-way ANOVA, followed by Dunnett’s multiple comparison tests. ****p < 0.0001, ns= p > 0.05.

Quantitative real-time PCR was used to investigate butein effect in *CCL2* gene expression in both breast cancer cell lines. The *CCL2* increased expression data had similar trend as the results the cytokine arrays and ELISA assays. TNF-α-induced *CCL2* expression was significant (p < 0.01) in both cell lines, compared to the control. Butein was effective in reducing *CCL2* expression significantly (p < 0.05) in MDA-MB-231 cells, causing inhibition of more than 50% in mRNA expression (**Figs 10A and B**). These results indicate that the changes in *CCL2* expression caused by butein at the transcriptome level, follow the same pattern observed at the protein level.

**Fig 10:**
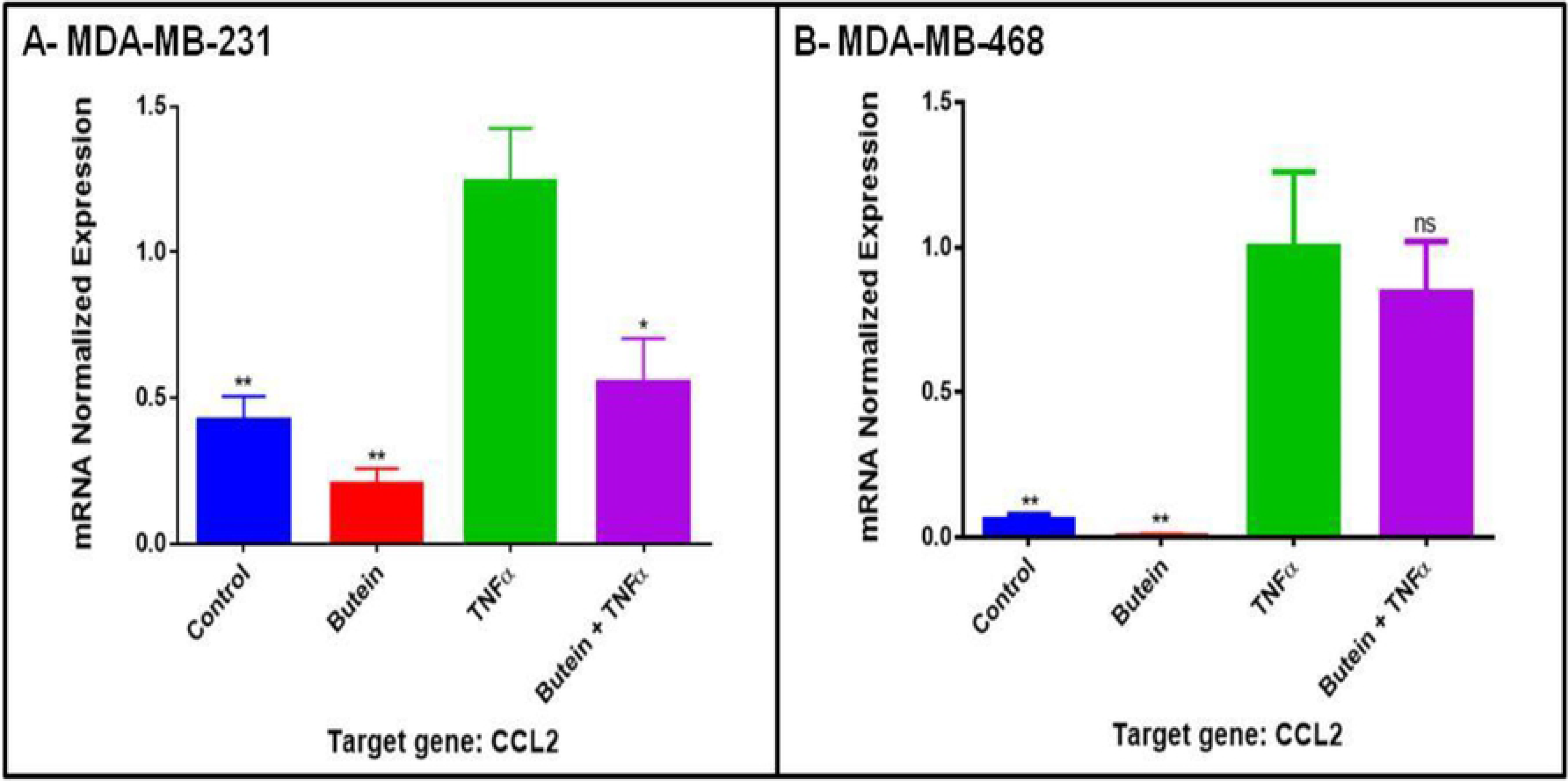
*CCL2* mRNA expression quantification in TNBC cells. The effect of butein (5 μM) in normalized *CCL2* mRNA expression in TNF-α stimulated MDA MB-231 (**A**) and MDA MB-468 cells (**B**). Each data point represents the mean ± S.E.M. of three independent studies (n=3). Statistically significant differences between TNF-α vs. other treatments were evaluated by a one-way ANOVA, followed by Dunnett’s multiple comparison tests. *p < 0.05, **p < 0.01, ns= p > 0.05.

To elucidate the possible signaling pathway related to the obtained findings, we investigated the changes in IKBKE mRNA expression. The results show that TNF-α upregulated IKBKE expression in both cells. TNF-α induced a 3.5 and 12.3-fold increase in mRNA expression in MDA-MB-231 and MDA-MB-468 cells, respectively, compared to the control. Although there was a higher expression of IKBKE in the TNF-α-stimulated MDA-MB-468 cells, butein co-treatment was only effective in MDA-MB-231 cells, inhibiting 37% of IKBKE mRNA expression (p < 0.05) (**Figs 11A and B**). These data demonstrate that IKBKE may be one of the NFκB signaling genes implicated in the TNF-α-induced *CCL2* release and its down-regulation by butein.

**Fig 11:**
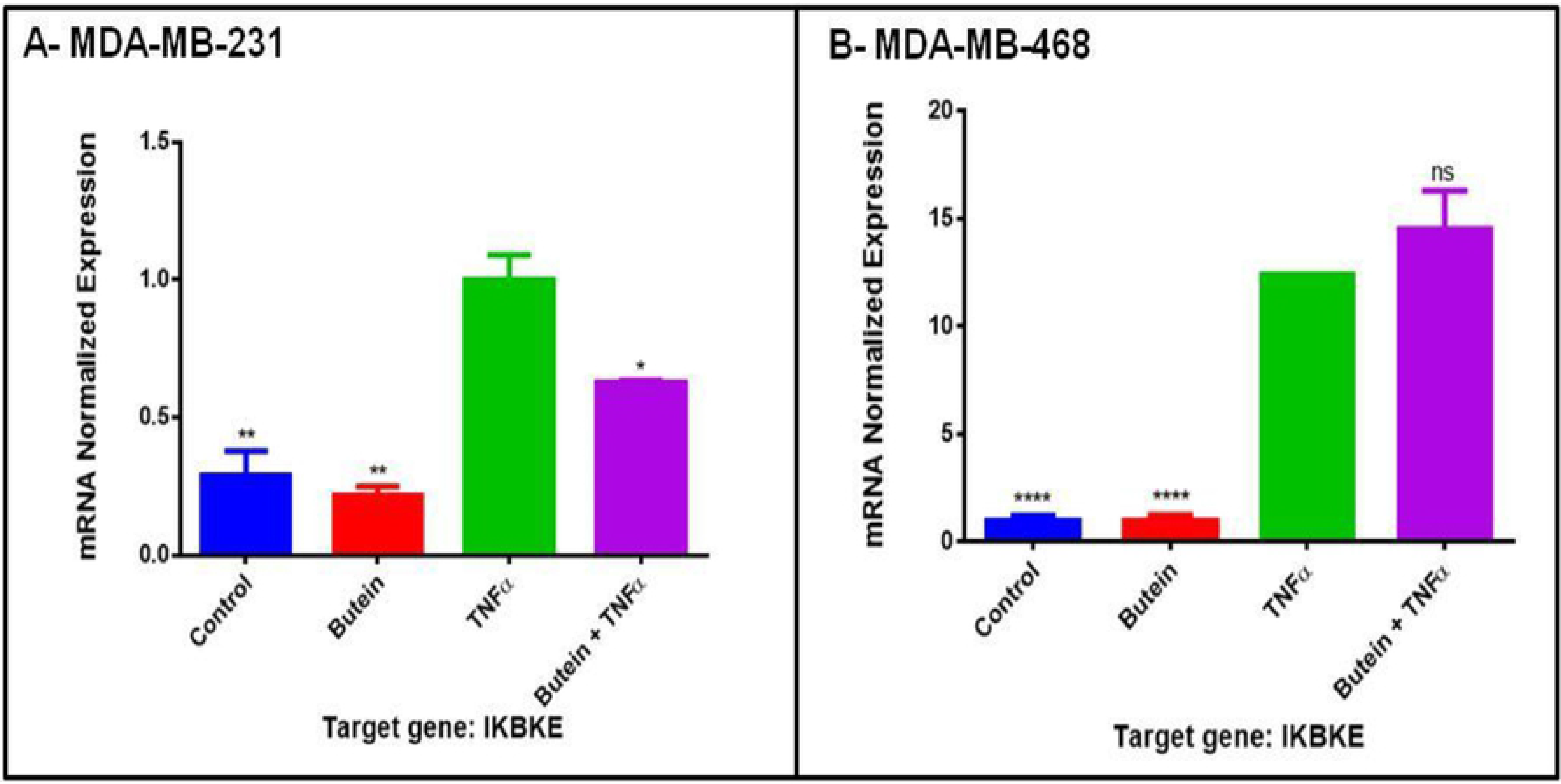
IKBKE mRNA expression in TNBC cells. The effect of butein (5 μM) in normalized IKBKE mRNA expression in TNF-α stimulated MDA MB-231 (**A**) and MDA MB-468 cells (**B**). The data points are expressed as the mean ± S.E.M. of three independent studies (n=3). Statistically significant differences between TNF-α vs. other treatments were evaluated by a one-way ANOVA, followed by Dunnett’s multiple comparison tests. *p < 0.05, **p < 0.01, ****p < 0.0001, ns= p > 0.05.

## Discussion

Polyphenolic compounds have received considerable attention for their use as a cancer chemopreventive and a chemotherapeutic agent. Previous *in vitro* studies showed butein cytotoxic and anti-proliferative effects on breast cancer cells, including MDA-MB-231 and MCF-7 [13,53], suggesting that butein might have similar effects in other breast cancer cell lines. However, there is no data comparing the effect of this compound in racially different TNBC cells. The current study shows butein anticancer properties in TNBC cells, specifically MDA-MB-231 and MDA-MB468, representing Caucasians and African Americans. Overall the results obtained in our study provide more evidence for butein cytotoxicity towards both cell lines. However, the compound highly impacted MDA-MB-468 cells, in which lower concentrations were more effective in reducing cell viability (**Figs 1, 2A and B**), and decreasing cell proliferation (**Fig 3**). These data corroborate with previous findings in the literature showing butein cytotoxic effects in MDA-MB-231 cells [13]. Also, the data show that butein induced apoptosis in both cell lines, increasing apoptotic cells ratio more effectively in MDA-MB-468 (**Figs 4A and B**; **5A and B**).

Proliferative and anti-apoptotic effects have been described to be associated with NFκB signaling activation, which induces cell growth and arrests programmed cell death in multiple cell lines [54–56]. NFκB activation was detected in ER negative breast cancer cell cultures [57], suggesting its role in proliferative pathways and cell death signals regulation [54–56]. In normal cells, activation of NFκB is an inducible and regulated event, where diverse stimuli, including proinflammatory cytokines (such as TNF-*α*), may initiate the proper stimulation and upregulate the transcription of many target genes. However, in tumor cells, impaired regulation of NFκB activation may lead to a deregulated expression of controlled genes [58]. Irregular NFκB activation can lead to many changes in the normal physiology of cells including self-induced growth signals, blockage in responses to growth inhibition, apoptosis evasion, persistent angiogenesis, invasion of tissues, and metastasis. NFκB can change cell homeostasis by inducing inflammatory processes, which have been described as associated with cancer development [59]. The association between cancer and inflammation has been extensively studied [60]. It is now clear that cell proliferation by itself doesn't cause cancer, however, the uncontrolled proliferation in an environment rich in inflammatory cells, DNA damage inducers, and growth factors, definitely potentiates and/or increases chances of tumor development [61].

Considering that the association of chronic inflammation with infection and irritation may promote environments that induce DNA lesions and tumor initiation [62], the current study investigated butein ability to inhibit TNF-α-mediated release of proinflammatory cytokines. The obtained findings in the current study show that butein attenuated the expression of CCL2, at both protein and mRNA levels in MDA-MB-231, but not in MDA-MB-468 cells (**Figs 8A and B**, **9A and B**, **10A and B**), demonstrating butein ability to inhibit CCL2 release only in Caucasians TNBC cells. CCL2 belongs to the C-C chemokines group and has been identified as an inflammatory modulator, which regulates macrophage recruitment during infection, the healing process, and autoimmune diseases. Through its CCR2 receptor affinity [63–65], it activates downstream signaling pathways, such as p42/44 MAPK, phospholipase C-γ, and protein kinase C. Elevated levels of CCL2 protein and mRNA expression are implicated in cancer, showing a high tumor grade and poor prognosis [66]. However, CCL2 inhibition in mammary tumor-bearing mice decreased tumor growth, metastasis, macrophage recruitment, and angiogenesis, suggesting that this cytokine regulates tumor progression via a macrophage dependent mechanism [50,67–71]. A previous study demonstrated that CCL2 treatment decreased apoptosis caused by serum deprivation, gentamicin or 5-FU treatment in mouse and human mammary carcinoma cells (MDA-MB-231), suggesting that CCL2 may induce pro survival effects in human breast cancer cells. Moreover, Fang *et al.* (2012) show that CCL2 effect on cell survival is linked to an increase of phosphorylation of Smad3 and p42/44 MAPK proteins [72].

The findings of our work demonstrate butein ability to induce apoptosis and inhibit TNF-α-induced CCL2 release, indicating an association between cell survival regulation and CCL2 inhibition. These data corroborate with previous literature studies showing the significant role of CCL2 signaling in breast cancer cells [73,74] and indicates that targeting CCL2 signaling pathway may affect various mechanisms involved in cancer progression, hence representing an attractive therapeutic target [72]. The present study also determined that butein inhibitory effect on CCL2 expression was only effective in MDA-MB-231 cells, suggesting that the apoptotic effect in MDA-MB-468 cells is not associated with CCL2 regulation.

Our investigation showed that butein inhibitory effect on CCL2 expression in Caucasian cells might be attributed to its ability to downregulate IKBKE mRNA expression (**Fig 11 A and B**). IKBKE is a gene overexpressed in approximately 30% of human breast tumors [75] and represents an emerging link between cancer and inflammation [76]. It promotes cytokine release and pro-survival signaling through the activation of NFκB and JAK–STAT signaling pathways [77]. Using JAK inhibitors that also target IKBKE, Barbie *et al.* [77], verified that there was a decrease in the viability of TNBC cells that presented *IKBKE* increased levels. This gene also regulates survival signaling associated with NFκB pathway activation, enabling cell transformation [76,78]. Also, Bauer *et al.* [79] studied the association between the *IKBKE* gene and CCL2 release, showing that *IKBKE* downregulation attenuates CCL2 expression in MDA-MB-231 TNBC cells. Likewise, the data from our previous study [80] demonstrated that the natural compound Plumbagin inhibited *IKBKE* gene expression and consequent release of CCL2 in TNF-α-induced MDA-MB-231 cells, strengthening the potential association between *IKBKE* and CCL2 expression.

In summary, the present investigation demonstrates butein potential in cancer suppression of two different TNBC cell lines: MDA-MB-231 and MDA-MB-468. Butein showed higher cytotoxicity, anti-proliferative, and apoptotic effects in MDA-MB-468, compared to MDA-MB-231. Additionally, butein downregulated both, protein and mRNA expression of TNF-α-stimulated CCL2 in Caucasian cells but not in African Americans. Moreover, the results elucidated one, out of many molecular mechanisms that may be involved in CCL2 downregulation, showing butein inhibitory effect on *IKBKE* mRNA expression in MDA-MB-231 cells (**Figs 12 and 13**). Therefore, the obtained findings indicate that butein might be a potential candidate in breast cancer therapy targeting CCL2 in Caucasians and may also provide an explanation regarding the poor response to therapy in African American patients with advance TNBC.

**Fig 12:**
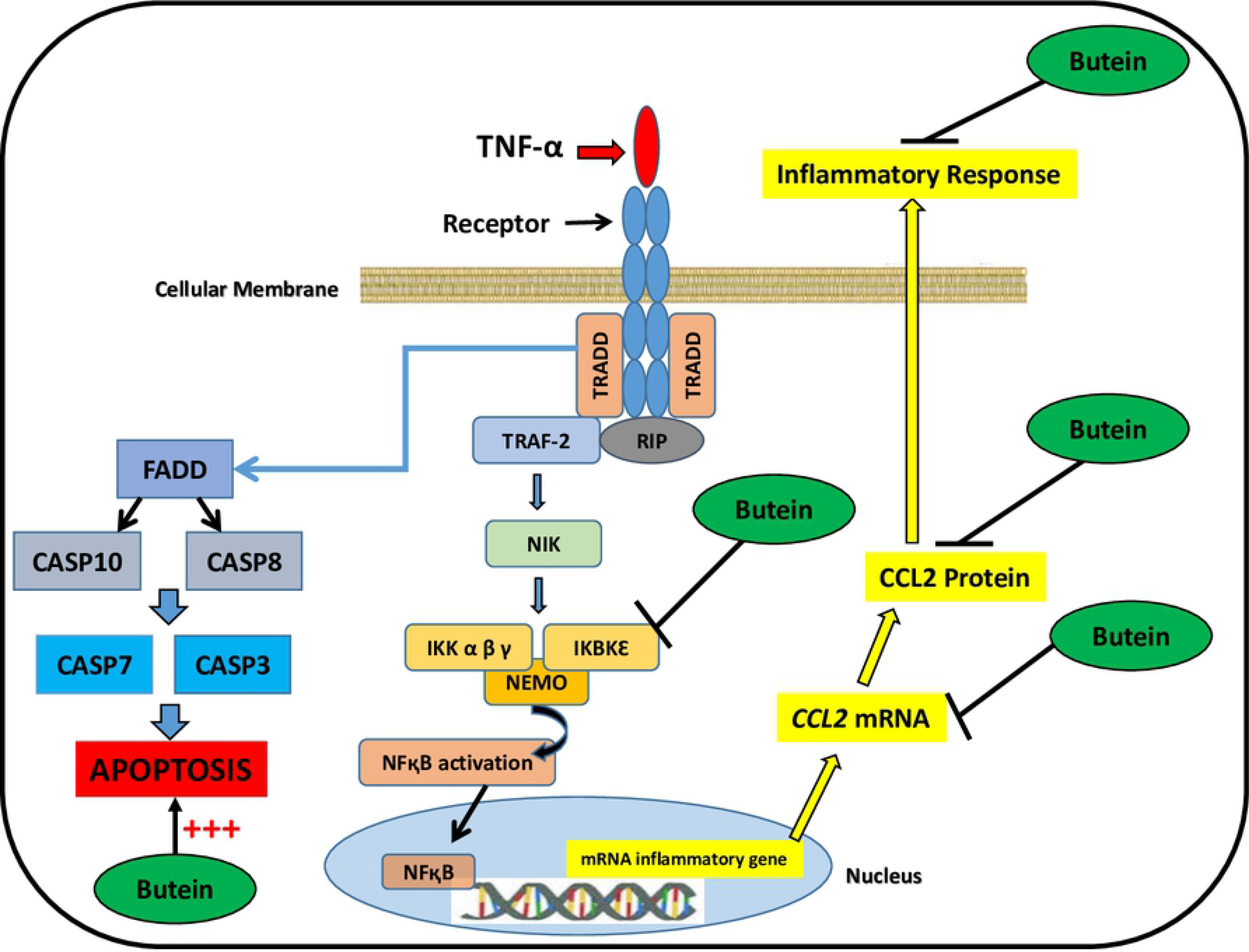
Proposed mechanism of butein effect in TNBC MDA-MB-231 cells. Diagram shows butein inhibitory effect in TNF-α-stimulated CCL2 expression at mRNA and protein level, attenuating *IKBKE* expression as a possible molecular mechanism, in addition to a possible signaling pathway for butein apoptotic effects in TNBC MDA-MB-231 cells.

**Fig 13:**
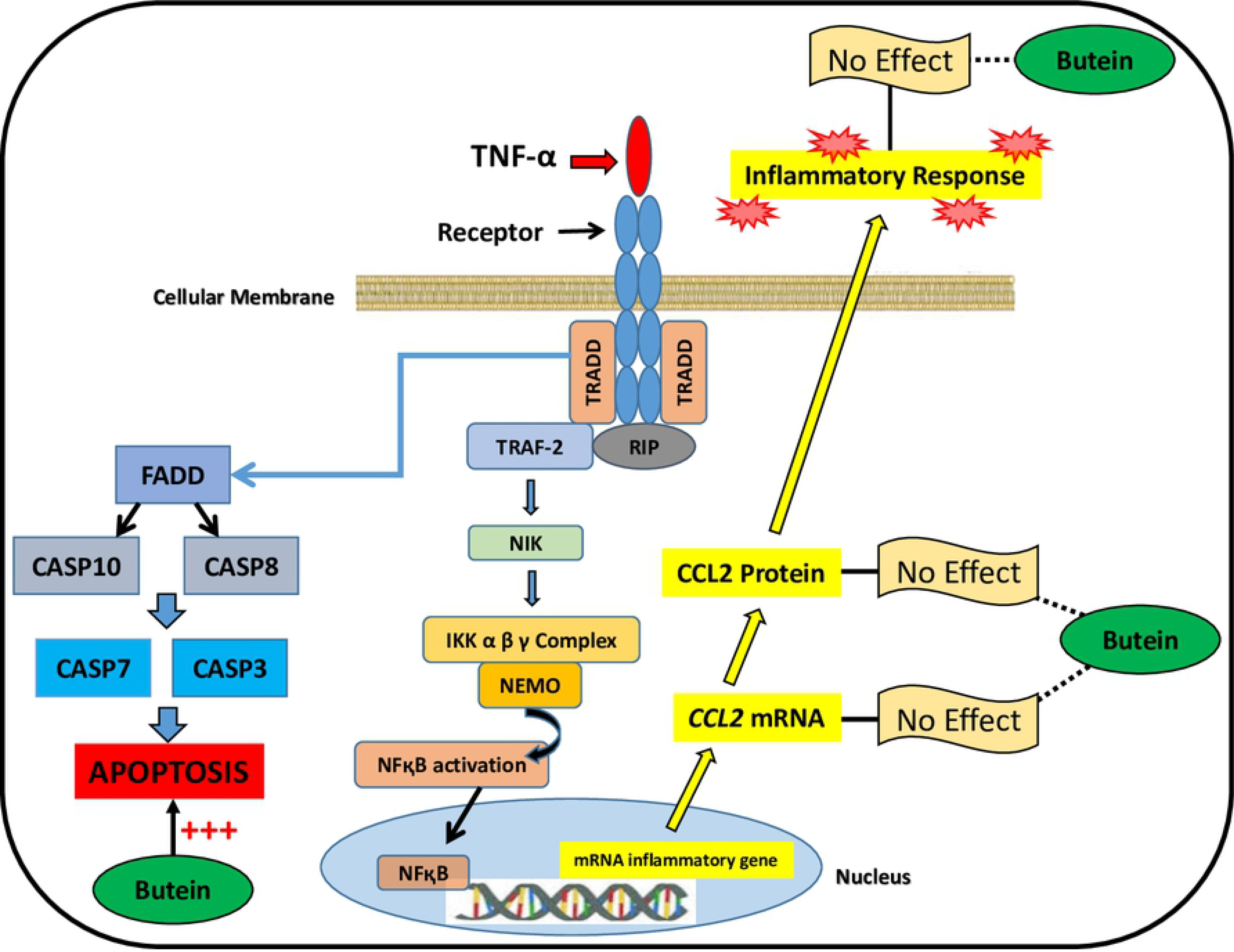
Proposed mechanism of butein effect in TNBC MDA-MB-468 cells. Diagram shows possible mechanism for butein apoptotic effects in MDA-MB-468 cells, and no significant effect in TNF-α-stimulated CCL2 expression.

## Authors' contributions

Conceptualization: PM, DB, KFAS

Methodology: PM, AH, DB, KFAS

Formal analysis: PM, AH, DB, SM, KFAS

Funding acquisition: KFAS

Project administration: KFAS

Resources: KFAS

Software: PM, SM

Supervision: KFAS

Writing original draft: PM

Writing review & editing: PM, AH, SM, KFAS

## Funding Statement

This work was supported by US National Institutes of Health (NIH), National Institute on Minority Health, and Health Disparities (NIMHD) grants G12 MD 007582 and P20 MD 006738 to KFAS.

## Data availability

All relevant data are within the manuscript.

